# Distinct heterozygous *TTC7A* missense variants lead to different intestinal epithelial phenotypes in pediatric IBD

**DOI:** 10.1101/2025.06.11.659065

**Authors:** Zahra Shojaei Jeshvaghani, Polina Deenichina, Lauren Collen, Maaike de Vries, Jesse Brunsveld, Daniel Kotlarz, Sibylle Koletzko, Christoph Klein, Jeffrey M. Beekman, Scott Snapper, Caroline A. Lindemans, Michal Mokry, Carmen Argmann, Ewart Kuijk, Edward Nieuwenhuis

**Affiliations:** Department of Pediatric Gastroenterology, Wilhelmina Children’s Hospital, University Medical Center, Utrecht University, 3584 EA, Utrecht, The Netherlands; Regenerative Medicine Utrecht, University Medical Center, Utrecht University, 3584 CT, Utrecht, The Netherlands; Division of Gastroenterology, Hepatology, and Nutrition, Department of Pediatrics, Boston Children’s Hospital and Harvard Medical School, Boston, MA 02115, USA; Department of Pediatric Pulmonology, Wilhelmina Children’s Hospital, University Medical Center, Utrecht University, 3584 EA, Utrecht, The Netherlands; Department of Pediatrics, Dr. von Hauner Children’s Hospital, LMU University Hospital, 80337, Munich, Germany; Department of Pediatrics, Gastroenterology and Nutrition, School of Medicine Collegium Medicum University of Warmia and Mazury, 11-082, Olsztyn, Poland; Division of Gastroenterology, Department of Medicine, Brigham & Women’s Hospital and Harvard Medical School, Boston, MA 02115, USA; Department of Stem Cell Transplantation, Princess Maximá Center for Pediatric Oncology, 3584 CS, Utrecht, The Netherlands; Laboratory of Experimental Cardiology, Department of Cardiology, University Medical Center Utrecht, Utrecht University, 3584 CX, Utrecht, The Netherlands; Laboratory of Clinical Chemistry and Haematology, University Medical Center Utrecht, 3584 CX, Utrecht, The Netherlands; Department of Genetics and Genomic Sciences, Icahn School of Medicine at Mount Sinai, New York, NY 10029, USA; Princess Maximá Center for Pediatric Oncology, 3584 CS, Utrecht, The Netherlands

**Author notes:** contributed equally (Ewart Kuijk, Edward Nieuwenhuis).

## Abstract

Pathogenic mutations in Tetratricopeptide repeat domain 7A (*TTC7A*) result in gastrointestinal and immunological disorders of which the pathobiology is not fully understood. Previous case reports indicate that TTC7A plays an important role in preserving intestinal epithelial integrity, but thus far only few variants have been investigated and it is unclear if different variants exert the same effects. Here, we aim to study the effects of different variants on the intestinal epithelium. We present three instances of pediatric inflammatory bowel disease (IBD), displaying varying clinical symptoms and severity levels, and associated with different heterozygous missense mutations in *TTC7A*. Intestinal organoids derived from patients show dissimilar epithelial phenotypes and exhibit differences in growth, morphology, apicobasal polarity, responses to specific drugs, TTC7A expression, and transcriptional profiles. The findings of our study suggest differences in pathobiology between individuals with different *TTC7A* mutations. This investigation enhances our comprehension of TTC7A-related conditions and can have implications for developing targeted therapies for TTC7A-associated disorders.

## Introduction

Mutations that interfere with the function of Tetratricopeptide repeat domain 7A (TTC7A) are associated with a range of intestinal and immune diseases with varying degrees of severity.^1–4^ To date, more than 90 patients with over 80 different *TTC7A* mutations have been reported.^5–7^ While missense mutations seem to cause very early onset inflammatory bowel disease (VEO-IBD), *TTC7A* nonsense mutations are suggested to be associated with more severe intestinal phenotypes like multiple intestinal atresia (MIA), which are accompanied by immunodeficiency in over 75% of cases.^1^ Several extraintestinal manifestations affecting the skin and/or hair have also been observed in patients with *TTC7A* mutations.^1,8–10^ Patients with complete TTC7A deficiency experience high rates of morbidity with a median survival age of less than 12 months.^1^ The most common causes of mortality in these individuals are typically sepsis, intestinal failure, pneumonitis, and complications related to hematopoietic stem cell transplantation (HSCT).^1,6^ Currently, there is no established standard of care for patients with *TTC7A* mutations. Despite the potential of HSCT to resolve immunologic problems, gastrointestinal issues persist, suggesting inherent intestinal epithelial dysfunction in patients with *TTC7A* mutations.^11–13^

Functional studies in human cell lines and patient-derived organoids indicate a role for TTC7A in maintaining intestinal epithelial homeostasis through multiple mechanisms. In HEK293T cells, wild-type TTC7A interacts with Phosphatidylinositol 4-kinase III alpha (PI4KIIIα) to synthesize plasma membrane PI4-phosphate (PI4P), essential for maintaining membrane identity, cell survival, and polarity.^14^ This process is deregulated by the IBD-associated *TTC7A* variants E71K, Q526X, and A832T.^15^ Consistently, *TTC7A* knockdown in Henle-407 cells reduces PI4KIIIα and PI4P levels.^15^ The importance of PI4P and PI4KIIIα for intestinal health is further supported by animal models.^16–18^ Small intestinal organoids derived from a patient with a homozygous nonsense variant (A832X) and in rectal organoids from another patient with a homozygous missense variant (E71K) exhibit aberrant activation of RhoA kinase (ROCK) effectors, causing epithelial defects in polarity, morphology, growth, and differentiation. Notably, Rock inhibitor treatment in these organoids reduced ROCK activation and restored the epithelial defects.^11,19^ Despite this, the exact mechanism of TTC7A involvement in the ROCK pathway remains unknown.^20,21^ Furthermore, TTC7A deficiency is associated with increased intestinal epithelial cell apoptosis, triggered by caspase cleavage.^11,15,19,22,23^ This observation has prompted the identification of leflunomide as a promising candidate for treating IBD resulting from TTC7A deficiency.^23^ Leflunomide is known to reduce inflammation by inhibiting pyrimidine synthesis in lymphocytes.^24^ However, via an alternative and unknown mechanism, in *TTC7A*-knockout cells, leflunomide effectively inhibited caspase activity by restoring AKT activation, an anti-apoptotic kinase.^23^ Moreover, leflunomide resolved multiple intestinal abnormalities in *ttc7a*-mutant zebrafish and patient-derived organoids with biallelic *TTC7A* mutations (E71K and L304R).^23^

Overall, despite evidence indicating possible functions of TTC7A and potential therapeutic targets for its deficiency, there remains a knowledge gap regarding the full extent of TTC7A’s roles and the potential impact of specific variants on epithelial cells in disease. Additionally, it is unclear to what extent candidate drugs such as ROCK inhibitors and leflunomide can be utilized to treat patients with varying *TTC7A* variants.

The objective of this study was to enhance comprehension of the impact of *TTC7A* mutations on intestinal epithelial phenotype and function. We identified distinct heterozygous missense *TTC7A* variants in three IBD patients and established long-term intestinal organoid cultures. These patient-derived organoids exhibited unique epithelial phenotypes and drug responses, suggesting that diverse molecular mechanisms underlie the disease across different *TTC7A* variants. Our results expand our knowledge of TTC7A pathobiology, potentially influencing future therapeutic efficacy for TTC7A-deficient patients.

## Material and Methods

### Research approval

Biopsy samples from clinically required surgical resections or diagnostic endoscopy, were collected from the patients and control subjects, along with clinical information, with their prior informed consent. Data collection and research studies were conducted in adherence to the ethical guidelines established by the ethics committees. Ileum biopsies from P1 and colon biopsies from P3 were acquired respectively from Boston Children Hospital (protocol # IRB-P00000529, titled: Pediatric Gastrointestinal Disease Biospecimen Repository and Data Registry) and Klinikum der Universitat Munchen (KUM protocol # 806-16, titled: Analysis of immunological and genetic causes in pediatric inflammatory bowel disease). Duodenal biopsies from P2, along with samples obtained from control subjects, were procured from the University Medical Center Utrecht (Medisch Ethische Toetsings Commissie, protocol # 10/402, titled: Specific Tissue Engineering in Medicine).

### Intestinal organoid cultures

Isolation of intestinal crypts from biopsy material was executed as previously described.^25^ In short, biopsies were washed with cold complete chelation solution and incubated in chelation solution supplemented with 10 mM EDTA for 30-60 minutes at 4 °C on a rocking platform. The supernatant was harvested, and EDTA was washed away. Crypts were isolated by centrifugation and plated in prewarmed 24-well plates, embedded in droplets of 50-70% Matrigel (growth factor reduced, phenol free; BD bioscience) diluted in basal medium (BM), consisting of advanced DMEM/F-12 (Gibco, 12-634-010), Penicillin-Streptomycin 50 U/mL (Gibco, 15070063), 10mM HEPES (Gibco, 15630080) and 1% GlutaMax (Gibco, 15630-056). Three times per week, organoids were refreshed with establishment medium composed of BM enriched with 0.5 nM Wnt-surrogate (U-Protein Express, N001-0.5mg), 2% Noggin conditioned medium (NCM) (U-Protein Express, N002-100 mL), 20% R-Spondin conditioned medium (derived from RSPO1-expressing HEK293T cells, kindly provided by Dr. C. J. Kuo, Department of Medicine, Stanford, CA), 50 ng/mL human EGF (Peprotech, 315-09-1MG), 10 mM Nicotinamide (SIGMA, N0636-500G), 1.25 mM N-acetyl cysteine (SIGMA, A9165), 1x B27 (Thermo Fisher, 12587010), 500 nM TGF-b inhibitor (A83-01, Tocris Bioscience, 2939/10), 10 μM P38 inhibitor (SB2021190, SIGMA, S7067-25mg), Primocin 100 μg/ml (InvivoGen, ant-pm-1). Once the organoid cultures were established, the organoid establishment medium was replaced with expansion medium (EM), which is identical except for the substitution of the Wnt surrogate and commercial NCM with 50% Wnt3a-conditioned medium and 10% NCM, respectively, both derived from in-house cell lines. For maintenance, organoids were passaged every 7-14 days, by mechanical dissociation in a ratio between 1:1 and 1:4, or by single cells using TrypleE (Thermo Fisher Scientific, 12604021). Single cell cultures were treated with EM supplemented with 10 µM ROCK inhibitor (Y-27632, Abcam, 120129) for the first 2-3 days. While TTC7A patient organoid cultures were supplemented with ROCK inhibitor for long-term maintenance, this was not applied during experimental procedures, unless otherwise specified. For the indicated experiments, leflunomide (Sigma, PHR1378-1G) was added to the EM to a final concentration of 10µM. All cultures were kept in incubators at 37°C, in a humidified atmosphere with 5% C0_2_. Morphology and overall condition of organoids was monitored using standard microscopy and the EVOS cell imaging system (Thermo Fisher Scientific). Analysis of organoid cellular phenotypes was conducted using PRISM software (Macintosh version 4, GraphPad Inc.).

### Organoid swelling assay

Methods to measure barrier integrity by means of forskolin induced swelling were adapted from protocols described previously.^26^ Briefly, mechanically dissociated organoids were seeded in flat-bottom 96-well culture plates in 4 μl of 50% Matrigel containing 30–50 organoids per well and immersed in 50 μl EM. One day later, organoids were stained with 3 µM calcein green (Invitrogen). Forskolin-induced swelling was monitored over a 1-hour period, capturing images at 7 time points with 10-minute intervals using confocal live cell microscopy (LSM710, Zeiss) at 37 °C. Organoid swelling was automatically quantified using Volocity imaging software (Improvision). The increase in total organoid area in the xy plane relative to the starting point (t = 0) under forskolin treatment was calculated and averaged based on data from two individual wells per condition. The area under the curve (AUC) (t = 60 minutes; baseline, 100%) was calculated using GraphPad Prism.

### Apoptosis assay

Caspase-dependent apoptosis in organoids was measured using the Caspase-Glo® 3/7 assay kit (Promega) according to the manufacturer’s protocol. Briefly, organoids were plated by seeding ∼1600 single cells per well in a 70% Matrigel droplet, in a 96-well culture plate. After one week, organoids were collected in 50 µL pre-cooled BM, transferred to a white 96-well plate, mixed with 50 µL Caspase-Glo® 3/7 reagent, and incubated at room temperature for 40 minutes. Luminescence was measured using a Tristar 3 Multimode microplate reader.

### Apicobasal polarity

Apicobasal polarity assay was carried out in 96-well clear flat bottom imaging plates. Following mechanical dissociation, ∼ 200 organoid fragments were seeded per well in a 50% Matrigel droplet and expanded for 3-4 days. For whole-mount immunostaining, organoids were fixed with 4% cold PFA, followed by permeabilization with 0.3% Triton X-100. Immunohistochemistry was performed as described previously ^14^ using the primary antibodies rat-anti-human CD49f binding integrin alpha 6 (2 µg/ml, BD Biosciences, 555734) and purified rat IgG2a isotype control (2 µg/ml, BD Biosciences, 555841), and the secondary antibody Goat-anti-Rat-IgG-(H+L)-Alexa 647-conjugate (2.5 µg/ml, Thermo Fisher, A21247). TRITC-labelled phalloidin (20 µg/ml, Sigma, P1951) was used to stain F-actin. DNA was stained with DAPI (5 µg/mL, Sigma, D9542). Images were acquired with a 20X objective on a Leica SPX8 confocal microscope controlled with the LAS X Software. Images were analyzed in ImageJ, and apicobasal polarity was assessed by visual determination of F-actin and integrin alpha 6 localization.

### Growth assay

10,000 single cells were cultured in 3 replicates for each condition in a 24-well plate. For the duration of 3 continual passages and before every mechanical splitting, the number of organoids in each condition (N=3) was counted manually using EVOS microscopy. The total number of organoids for each condition in one passage, was obtained by multiplying the number of counted organoids in that passage by the previous splitting ratios.

### Protein blotting

Intestinal organoids were isolated from Matrigel using cold BM, washed with cold PBS for 2 times, and lysed in Laemmli Lysis Buffer (LLB) (4% SDS, 20% Glycerol, 120 mM Tris-HCL pH 6.8) supplemented with protease inhibitors (Roche). Protein concentration was measured following the Thermo Fisher BCA kit protocol. A 20x Sample Buffer (50% LLB, 0.1% Bromophenol Blue, 2.5% β-Mercaptoethanol) was diluted into 50 µg of protein lysate in a total volume of 50 µL. Samples were heated at 95°C for 5 minutes, separated by SDS-PAGE, blotted, and then stained with the specific monoclonal antibodies against TTC7A (1 µg/ml, Origene, TA812302) and β-actin (40ng/ml, Santa Cruz, sc-47778). After staining with a horseradish peroxidase-conjugated secondary antibody, the immunoblot was developed with an enhanced chemiluminescence detection kit (Thermo Fisher’s SuperSignal West Dura Extended Duration Substrate). Signal intensities were visualized in a ChemiDoc Touch Imaging System (Bio-Rad).

### RNA sequencing analysis

From a previously established RNA-seq dataset of intestinal organoids, which included samples at baseline and after immunological stimulation, we selected and analyzed the baseline samples (n=61). This cohort comprised samples from 17 unique non-IBD donors and 23 unique pediatric IBD patients, including three TTC7A patients reported in this study **(Supplementary Tables 1 and 2)**. The dataset included duplicate samples and samples from multiple intestinal regions (colon and ileum).

Raw count data was pre-filtered to keep genes with CPM>1.0 for at least 30% of the samples (n=13625 genes). After filtering, count data was normalized via the weighted trimmed mean of M-values.^27^ Gene expression matrices were generated using the voom transformation^28^ and adjusted for technical variables (e.g. batch and organoid intestinal location) using the limma framework **(See QC plots in Supplementary Figure 1)**. Multiple biological samples per subject were accounted for using duplicateCorrelation function in R. Statistical analysis to determine genes differentially expressed in each of the three TTC7A patient samples vs healthy control (HC) was carried out using the limma framework in R language version 4.04^29^ and its available packages using the corrected expression matrix. Differentially expressed genes (DEGs) were defined according to p value<0.05. This cut-off was used (instead of Adj p value) as technical replicates for the TTC7A individual patients were limited in number (N=2).

Weighted gene coexpression network analysis (WGCNA)^30^ was performed in R using adjusted lcpm transformed count data (n=61 samples, 13625 genes) essentially as previously described.^31^ A signed network (beta = 12) was generated using blockwiseModules function (maxBlockSize = 30000) with default parameters except minModuleSize=50. See **Supplementary Figure 2** for scale free plots and clustering analysis. Pathway enrichment analysis for DEGs or WGCNA modules was performed using Fisher’s exact test in R with Benjamini Hochberg (BH) multiple test correction using genesets sourced from Enrichr ^32^ and included Gene Ontology Biological process (GO BP, 2023) and Reactome (2019). Packages available in R were used to generate the Upset plot ^33^ and heatmaps (gplots).

## Results

### Clinical features of patients

#### Patient 1

Patient 1 (P1), an 8-year-old male at the time of biopsy collection, was firstly diagnosed with neuropathic chronic intestinal pseudo-obstruction (CIPO) and microcolon following an exploratory laparotomy in the neonatal period. As a result of the associated intestinal failure secondary to CIPO, P1 has been dependent on ileostomy to receive parenteral nutrition throughout his life since infancy. At approximately 9 years old, P1 developed Crohn’s-like disease. Endoscopic evaluations over the next 10 years consistently demonstrated duodenal and ileal inflammation, which ranged in severity from mild to severe. This condition led to symptoms such as hematochezia (passing of blood through the rectum) and anemia. An ileal stricture was identified at one point and ultimately resected at age 17. P1 also had left-sided colonic inflammation with two associated strictures in the transversum and rectosigmoid. Notably, even though P1 did not experience frequent infections or exhibit clinical features indicative of combined immunodeficiency (CID), this patient showed IgA deficiency and lymphopenia affecting T and B cells. Medical history also includes short stature (height z-score −1.50), steroid-induced diabetes mellitus, delayed puberty, food allergies, and seborrheic dermatitis.

Genetic analysis of P1 revealed two deleterious known *TTC7A* variants in Exon 16 (c.1817 A>G; p.K606R) and Exon 17 (c.2014T>C; p.S672P) on the paternal allele as well as a *TTC7B* heterozygous mutation of unknown significance (c.2224 C>A; p.L742I) on the maternal allele.^4^

Throughout this course, P1 has been treated with aminosalicylates, anti-TNF, thalidomide, octreotide, and steroids, without control of the bleeding.

#### Patient2

Patient 2 (P2), a 13-year-old male at the time of biopsy collection, was diagnosed with mild colitis and CID when he was 4 years old. His intestinal condition was initially misdiagnosed with microvillous inclusion disease (MVID) based on histology. The same was true for his little brother who had a more severe intestinal disease and CID. Additionally, in P2, liver enzymes were found to be elevated, and the ultrasound showed liver abnormalities, consistent with previously reported PN-unrelated hepatic complications in other TTC7A patients.^9,11,19^ P2 experienced prolonged recovery periods from gastrointestinal infections.

At the age of 12, P2 was identified with compound heterozygous *TTC7A* mutations including, a novel mutation in Exon 11 (c. 1355 T>C; p.L452P) and a mutation in Exon 4 (c. 518G>T; p.G173V) that was previously reported by our group.^7^ This same combination was also found in his brother, who had at that time already passed away from the complications of HSCT.

P2 underwent HSCT with a cord blood donor to treat his TTC7A immunodeficiency. Following the transplantation, he experienced mild Graft-versus-Host Disease (GvHD) of liver and intestine, from which he eventually recovered. Throughout the disease course, P2 has been treated with immunosuppressant (Neoral), corticosteroid, omeprazole, granisetron, cidofovir, voriconazole, immunoglobulin therapy (Nanogam), amlodipine, tramadol and paracetamol. For years after the transplant, in case of fever or symptoms of diarrhea, P2 received prophylactic antibiotics. Otherwise, P2 is currently in good health and has discontinued the use of immunosuppressants within 6-12 months post HSCT.

#### Patient 3

Patient 3 (P3), a 14-year-old male at the time of biopsy collection, was diagnosed with chronic watery diarrhea, vomiting and intestinal insufficiency requiring total parenteral nutrition, and CID in the first year of age. The condition was accompanied by severe pyloric and jejunal stenosis and multiple stenosis requiring regular balloon dilatations. He showed active gastritis with eosinophilic infiltrates and high apoptosis with squamous (esophageal) metaplasia and from 18 years onwards also intestinal metaplasia in the corpus. To meet nutritional needs, P3 received a combination of home parenteral nutrition and minimal enteral feeding through a jejunostomy tube until the age of 14 years, alongside small amounts of oral intake. P3 has had a history of recurrent fever and multiple infections, including *Norovirus* infection, Enterococcal septicemia, *Salmonella* infection of the gallbladder, cytomegalovirus (CMV) infection affecting pulmonary and gastrointestinal systems, and herpes zoster ophthalmicus. This patient has also had other documented extraintestinal complications, including Acute Respiratory Distress Syndrome (ARDS), allergic rhinoconjunctivitis, food allergies, and a growth hormone secretion disorder.

Genetic analysis of P3 identified compound heterozygous *TTC7A* mutations in Exon 8 (c. 1027 G>A; p. E343K) and Exon 20 (c. 2405 T>C; p. L802P), previously reported by our group an others.^5,7^ Furthermore, another heterozygous mutation in Factor V Leiden was identified in this patient.

Prior to genetic diagnosis, at the age of 5 years, P3 received an allogenic HSCT from his HLA-identical but blood group discordant older brother, complicated by mild hepatic veno occlusive disease (VOD) and GvHD of the liver. Although his CID was corrected (normal immunoglobulins and no further serious infections), his gastrointestinal manifestations improved only slightly.

The clinical overview of the three patients is summarized in **Table 1**, representing how diverse *TTC7A* variants result in a range of heterogeneous clinical manifestations, varying in severity and necessitating distinct medical interventions for each of these individuals. We assessed the pathogenicity of the identified patient *TTC7A* missense variants using PolyPhen-2, SIFT, MutationTaster 2021, and AlphaMissense.^34–37^ While there was consensus among these prediction tools on the pathogenic nature of G173V (P2), L452P (P2) and L802P (P3), MutationTaster 2021 and AlphaMissense differed from PolyPhen-2 and SIFT in their predicted impact for K606R (P1), S672P (P1) and E343K (P3) **(Table 2)**. This highlights the limitations of computational prediction tools and underscores the necessity for functional evaluation to confirm the pathogenicity of these *TTC7A* variants. In our previous study, we functionally assessed K606R (P1), G173V (P2), E343K (P3), and L802P (P3) in Caco-2 cells. K606R and G173V were identified as pathogenic based on their induction of aberrant transcriptional signatures, whereas E343K and L802P were considered benign or inconclusive due to their limited impact on epithelial phenotype.^7^

**Table 1.** Patient History and Clinical Overview.

| Patients |  | P1 | P2 | P3 |
| --- | --- | --- | --- | --- |
| Age at IBD Diagnosis (years) |  | 9 | 4 | 1 |
| Gender |  | M | M | M |
| IBD Manifestation |  | Crohn's like | Mild colitis | Active eosinophilic gastric with squamous cell and intestinal metaplasia and atrophic |
| enteropathy |  |  |  |  |
| Immune deficiency |  | IgA deficiency, lymphopenia | CID | CID |
| Other Diagnoses |  | Neuropathic CIPO, microcolon | GvHD after HSCT | Pyloric and upper jejunal stenosis |
| Extraintestinal Conditions |  | Food allergies, seborrheic dermatitis, steroid-induced diabetes mellitus, delayed puberty, short stature | Hepatic abnormalities and GvHD after HSCT | ARDS, Hepatic VOD and GvHD after HSCT, GHD, multiple food allergies |
| TTC7A Mutation |  | H. c.1817A>G; p.K606R<br>c.2014T>C; p.S672P | CH. c.518G>T; p.G173V<br>c.1355T>C; p.L452P | CH. c.1027G>A; p.E343K<br>c.2405T>C; p.L802P |
| Intestinal Stricture |  | Yes | None | Yes |
| HSCT |  | No | Yes | Yes |
| Surgical History |  | Exploratory laparotomy, ileostomy | None | Multiple balloon dilatations of the pylorus, jejunostomy, cholecystectomy |
| Nutritional Source |  | TPN | Oral nutrition | HPN since early infancy, partly oral nutrition |

IBD, Inflammatory bowel disease; CIPO, Chronic Intestinal Pseudo-Obstruction; CID, Combined Immunodeficiency; GvHD, Graft-versus-Host Disease; ARDS, Acute Respiratory Distress Syndrome; VOD, Veno-occlusive disease; GHD, Growth Hormone Deficiency, H., Heterozygous; CH., Compound Heterozygous; SCT, Stem Cell Transplantation; BMT, Bone Marrow Transplantation; TPN, Total Parenteral Nutrition; HPN, Home Parenteral Nutrition.

**Table 2.** Predicted Pathogenicity of Patient *TTC7A* Variants.

| Patient | Variant | PolyPhen2 | SIFT | MutationTaster<br>2021 | AlphaMissense |
| --- | --- | --- | --- | --- | --- |
| **P1** | **K606R** | Pathogenic | Pathogenic | Benign | Benign |
|  | **S672P** | Pathogenic | Pathogenic | Benign | Benign |
| **P2** | **G173V** | Pathogenic | Pathogenic | Pathogenic | Pathogenic |
|  | **L452P** | Pathogenic | Pathogenic | Pathogenic | Pathogenic |
| **P3** | **E343K** | Pathogenic | Pathogenic | Benign | Pathogenic |
|  | **L802P** | Pathogenic | Pathogenic | Pathogenic | Pathogenic |

We hypothesized that these missense mutations result in hypomorphic variants, leading to reduced protein activity. Such variants are often associated with diverse clinical outcomes and severity, depending on the degree of gene impairment caused by the hypomorphic mutation. As seen in our cases, in the instance of P1, the intestinal ailment displays more severe symptoms compared to the immune disorder, characterized primarily by IgA deficiency and lymphopenia, which may not be directly attributable to the identified *TTC7A* mutations. On the other hand, for P2, the immune deficiency indicated by CID is prioritized over the mild intestinal colitis, explaining the effectiveness of HSCT in this patient. For P3, both the intestinal and immunological disorders hold significant clinical importance, with intestinal strictures and CID being showcased respectively. This severe disease phenotype typically diverges from what is commonly associated with hypomorphic variants of *TTC7A.*^1^

### Intestinal epithelial organoids from patients with different TTC7A mutations show different phenotypes

TTC7A patient-derived intestinal organoids exhibit epithelial defects in growth, morphology, viability, and polarity.^11,19,23^ To investigate the connection between different *TTC7A* variants and their effect on the linked epithelial impairments and responses to drugs, we generated organoids from biopsies taken from the ileum, duodenum, and colon of P1, P2, and P3, respectively. We then compared each patient-derived organoid line with organoid lines from healthy donors that originated from corresponding anatomical locations. Unlike organoids from P1 and P2, those from P3 exhibited reduced growth and poor recovery after freezing and thawing, even when treated with ROCK inhibitor. As a result, our investigation on P3 organoids was limited to morphology and TTC7A expression.

#### Growth, morphology, apoptosis and polarity

In the presence of ROCK inhibitor, organoids from P1 and P2 followed a normal growth pattern similar to that of control organoid lines from healthy donors (weekly expansion ratio of 1:2-1:4). Colon organoids from P3 grew more slowly with a ratio of 1:2 or less and eventually ceased to proliferate after six passages post-thaw **(Figure 1A)**.

**Figure 1.**
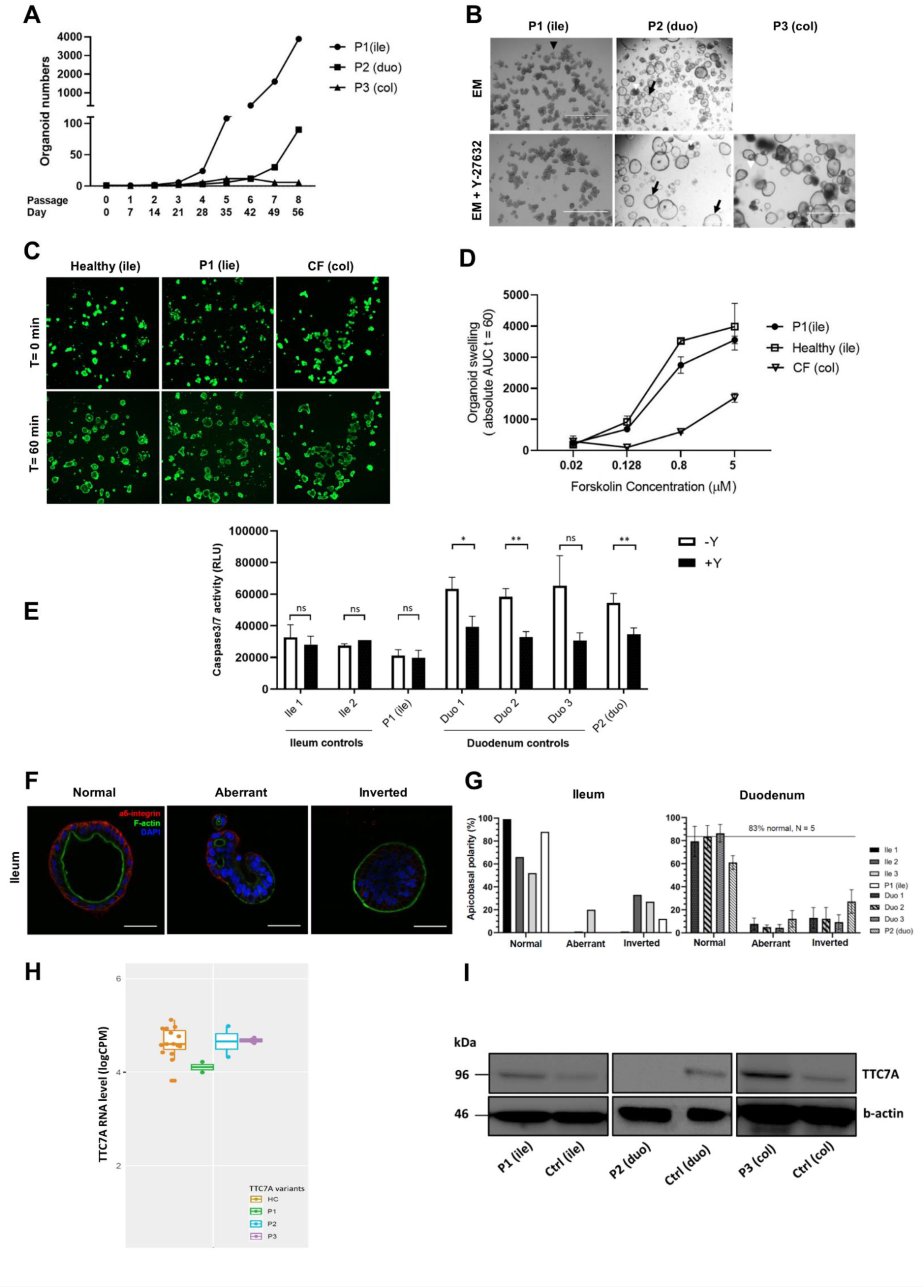
Intestinal organoids from patients with distinct *TTC7A* mutations show different epithelial phenotypes. **(A)** Growth profiles of patient-derived intestinal epithelial organoid cultures over continual passages. P1 (ile), P2 (duo) and P3 (col) represent organoid cultures originated from ileum, duodenum and colon biopsies of three patients with different mutations in *TTC7A*. **(B)** Representative bright field images of patient-derived intestinal organoid cultures in expansion medium in the absence (EM) or presence (EM+Y27632) of ROCK inhibitor (10µM). Different organoid morphologies are presented, including cystic structures (black arrow) with swollen, circular organoids, collapsed structures (black arrowhead), characterized by densely packed, non-cystic formations and aggregated structures (white arrowhead) where distinct shapes could not be identified. Images were obtained with a stereomicroscope (EVOS) at a magnification of 4x. **(C)** Representative confocal microscopy images of calcein green–labeled control Healthy (ile), TTC7A P1 (ile) and cystic fibrosis colorectal, CF (col), organoids with F508del/S1251N CFTR mutations as the negative control before and after 60 minutes of stimulation with forskolin (5µM). Scale bars, 200 μm.**(D)** Quantification of swelling in organoids (N = 2) derived from control Healthy (ile), TTC7A P1 (ile) and CF (col) after 60 minutes of stimulation with different concentrations of forskolin, calculated as the area under the curve (AUC) (Mean ± SD). **(E)** Caspase 3/7 activity in organoid cultures in the absence (-Y) or presence (+Y) of ROCK inhibitor, measured 7 days after seeding 1600 cells/well of a 96-well plate. Ile 1-2 (ileum) and Duo 1-3 (duodenum) are healthy control lines for P1 (ile) and P2 (duo), respectively. (Mean + SD, N = 3,*p </= .05, **p </= .01, multiple unpaired *t* test). **(F)** Representative immunofluorescent images of organoids with Normal, aberrant and inverted apicobasal polarity. Normal polarity is characterized by expression of α6-Integrin at the basolateral, thus outer membrane, F-actin ring that shows apical expression and lines the lumen of the organoids. Inverted polarity is characterized by F-actin expression at the entire outer membrane of the organoids and α6-Integrin inside the F-actin ring. Aberrant polarity is assigned to organoids that show expression of both α6-Integrin and F-actin at their outer border, to organoids with multiple F-actin lined lumens and also to organoids that completely lack the expression of either α6-Integrin or F-actin. Magnification 63x. Scale bar, 50 µm. **(G)** Quantification of organoids with normal, aberrant, and inverted polarity in 4-6 wells of a 96-well plate, 4 days after plating 100-200 organoid fragments per well. Healthy control lines of ileum (ile 1-3) and duodenum (duo 1-3) for the respective P1 (ile) and P2 (duo) TTC7A-deficient patient-derived organoids (Mean + SD). **(H)** *TTC7A* RNA expression in organoids. Count matrix was adjusted for batch and location and the resulting logcpm of TTC7A plotted and colored by disease status with HC = healthy control and either P1, P2 or P3 *TTC7A* variant patients (N=2 for each patient line). **(I)** Western blot analysis of TTC7A protein expression in organoids derived from TTC7A patients and their respective controls. b-actin was used as a loading control.

Intestinal organoids from the three TTC7A patients exhibited notable differences in appearance. When compared to healthy organoids from similar anatomical regions of the intestine, organoids derived from P1 displayed a completely collapsed appearance without a central lumen, while those from P2 generally appeared in a normal cystic phenotype **(Figure 1B and supplementary Figure 3A and B)**. Organoids derived from P3 displayed as cystic structures, along with the presence of aggregated formations where distinct shapes were indiscernible **(Figure 1B and supplementary Figure 3A)**.

To determine whether the collapsed morphology of organoids in P1 results from impaired epithelial barrier function, we measured organoid swelling potential, a method previously employed in a study to assess epithelial functionality in colon organoids from a patient with biallelic *TTC7A* mutations (E71K and L304R).^23^ The addition of forskolin induces fluid secretion by increasing intracellular cyclic AMP (cAMP) levels, which in turn activates anion and fluid transport into the organoid lumen mediated by the cystic fibrosis transmembrane conductance regulator (CFTR). The swelling of P1 organoids caused by forskolin-induced fluid secretion was similar to that of healthy control organoids **(Figure 1C and D)**, demonstrating that epithelial ion transport function and barrier integrity are unaffected in P1 organoids, which is in contrast to previous findings on colon organoids of a patient with *TTC7A* mutations.^23^

TTC7A deficiency has been associated with increased caspase-dependent apoptosis in intestinal epithelial organoids^19^ and epithelial cell lines.^15,23^ However, P1 and P2 organoids showed no increase in caspases 3 and 7 activity compared to controls, regardless of ROCK inhibitor treatment **(Figure 1E)**.

*TTC7A* mutations have also been linked to polarity defects.^11,15,19^ To investigate polarity, we performed immunofluorescence for F-actin and integrin alpha 6, to respectively mark the apical and basolateral membranes **(Figure 1F)**. P1 did not show polarity defects when compared to controls. In contrast, approximately 35% of all investigated P2 organoids displayed abnormal phenotypes, with roughly two-thirds being inverted and one-third being aberrant **(Figure 1G)**.

#### TTC7A expression

RNA sequencing results showed no differential expression of *TTC7A* between the patients or when compared to healthy organoid lines **(Figure 1H).** However, western blot analysis revealed variation in TTC7A expression at the protein level among the organoid lines derived from our patients. While P1 and P3 organoids expressed TTC7A protein at levels comparable to healthy controls, no TTC7A expression was detected in P2 organoids **(Figure 1I and supplementary Figure 4)**, indicating reduced TTC7A protein translation or stability in P2 organoids.

Consistently, in our previous study using TTC7A^-/-^ Caco cells, none of the relevant missense mutations (K606R in P1, G173V in P2, E343K in P3, and L802P in P3) altered TTC7A expression at either the RNA or protein level compared to the WT variant. This aligns with our current findings in patient-derived organoids, except for P2 organoids, which show reduced TTC7A protein expression.^7^

#### Response to Drugs

The RhoA-ROCK signaling pathway is recognized for its role in regulating the assembly of the actin cytoskeleton.^38^ *TTC7A* mutations are associated with improper activation of ROCK pathway in epithelial organoids.^19^ Accordingly, ROCK inhibition with Y-27632 reversed actin-related epithelial abnormalities such as polarity and morphology and also restored proliferation and normal culture expansion in ileal organoids from a patient with homozygous nonsense mutations (A832X)^19^ and in rectal organoids from another patient with homozygous missense variants (E71K).^11^ Here, we explored if ROCK inhibition would have similar beneficial effects on P1 and P2 organoids. While Y-27632 corrected the inverted polarity, it did not have a significant effect on the number of organoids with aberrant polarization, including those with multiple lumens, lacking the expression of either integrin alpha 6 or F-actin, or expressing both markers at the basolateral membrane **(Figure 2A)**. This suggests that the aberrant phenotypes of polarization in P2 organoids is not mainly driven by the activated ROCK pathway. Addition of Y-27632 did not lead to a cystic morphology of the organoids from P1 **(Figure 1B)**, implying the collapsed morphology is not the sole consequence of an enhanced RhoA kinase activity. However, Y-27632 significantly increased cystic morphology in both healthy and P2 duodenal lines **(Figures 1B and 2B)**. While ROCK inhibitor was dispensable for P1 organoid growth and had no impact on culture expansion or caspase-dependent apoptosis in P1 and ileal controls **(Figures 1E and 2C)**, it significantly enhanced the expansion of P2 and healthy duodenal organoids **(Figure 2D)**. This enhanced growth in duodenal organoids is likely due to reduced apoptosis in the presence of ROCK inhibitor **(Figure 1E)**, facilitating long-term culture regardless of *TTC7A* mutations.

**Figure 2.**
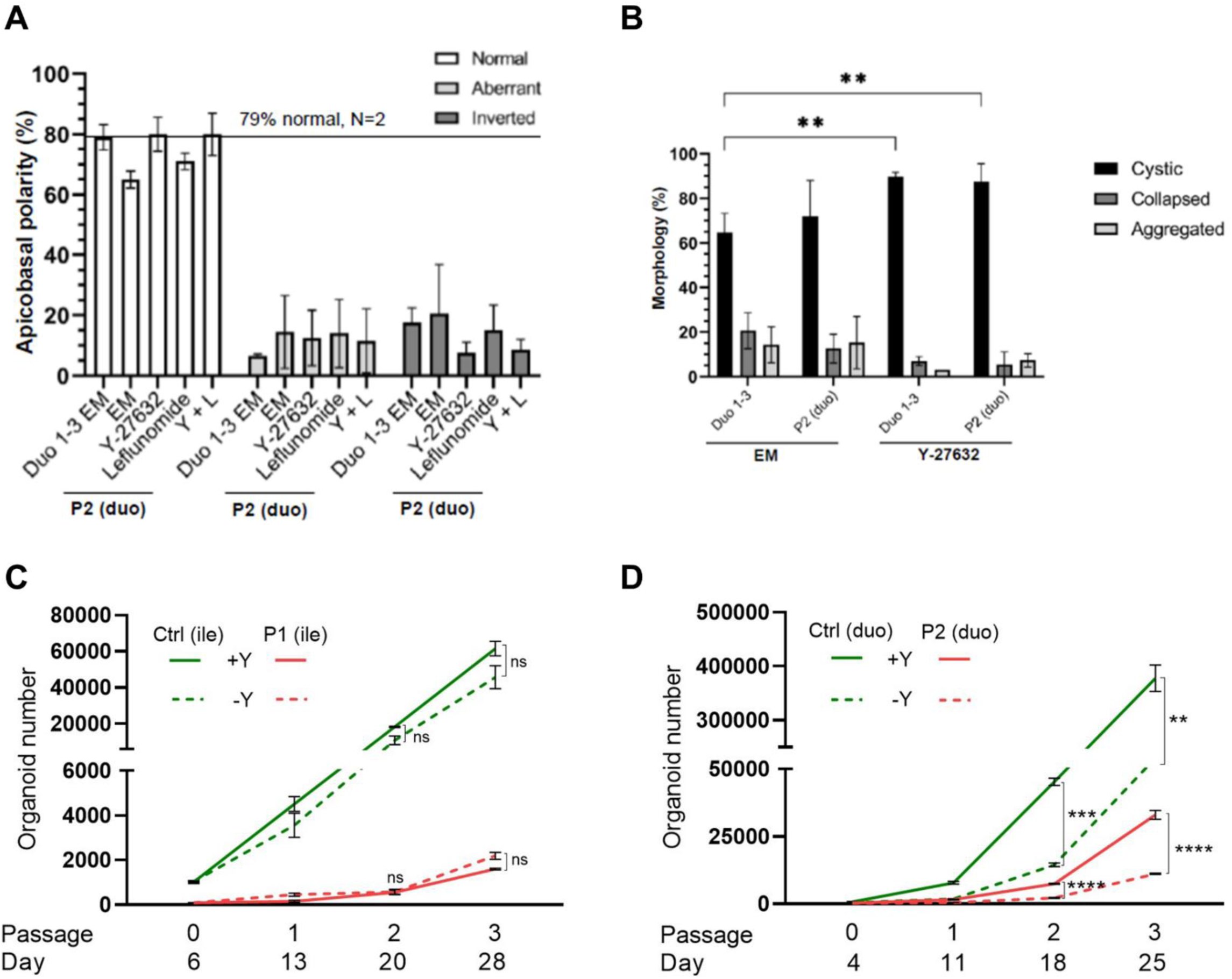
Intestinal organoids from patients with distinct *TTC7A* mutations show different responses to drugs. **(A)** Quantification of apicobasal polarity in 4-8 wells of a 96-well plate, 4 days after plating100-200 organoids/well treated with expansion medium without drug (EM) or supplemented with ROCK inhibitor (Y-27632), Leflunomide or both (Y+L) (10µM each), N=2. **(B)** Morphological comparison of duodenum organoids between groups treated with (Y-27632) and without (EM) ROCK inhibitor. Morphology was assigned as cystic, collapsed, or aggregated by counting 50 organoids per field (N=3) in ImageJ. Values were normalized to the total number of organoids for each group and averages of each experiment were used to perform 2-way ANOVA statistical analyses (Mean ± SD, *p </= .05, **p </= .01, ***p </= .001, ****p </= .0001). **(C)** Growth profiles for organoids derived from P1 (ile) and **(D)** P2 (duo) compared to their respective healthy control organoid lines in the presence or absence of Y-27632. Mean organoid number after passaging of 3 independent wells each. (Mean ± SEM, *p </= .05, **p </= .01, ***p </= .001, ****p </= .0001, multiple unpaired *t* test).

Leflunomide is an immunomodulatory drug that inhibits dihydroorotate dehydrogenase (DHODH), thereby reducing pyrimidine synthesis in lymphocytes.^24^ As a potential treatment for TTC7A-associated intestinal disease, leflunomide has been described to partially rescue cytoskeletal and polarity defects in colon organoids from a patient carrying compound heterozygous *TTC7A* missense mutations (E71K and L304R).^23^ This effect appears to be exerted through an alternative, yet uncharacterized, mechanism of action whereby it increases activated AKT and promotes cell survival.^23^ In P2 organoids, however, leflunomide failed to restore the polarity to homeostatic levels. Furthermore, the combination of leflunomide with ROCK inhibition offered no improvement over ROCK inhibition alone **(Figure 2A)**.

Taken together, our findings reveal a significant phenotypic heterogeneity among patients with distinct *TTC7A* mutations in their organoids. Variations in growth, morphology, polarity, and TTC7A protein levels were observed amongst patient organoids. It is worth noting that in contrast to previous research conducted on organoids with different *TTC7A* mutations,^11,19,23^ disruption of polarity or impaired epithelial transport function was not detected in P1. Elevated caspase activity was also not observed in P1 or P2. Furthermore, organoids exhibited different responses to ROCK inhibition or treatment with leflunomide.

### Diverse transcriptional profiles amongst patients with differing TTC7A mutations

To identify the molecular differences that may underlie the phenotypic variation amongst the three patients with *TTC7A* variants, we performed RNA-seq analysis on cultures of their intestinal epithelial organoids and compared the expression profiles to that of healthy controls. Given the small sample sizes associated with rare patient material, we considered genes differentially expressed using an unadjusted P <0.05 **(Supplementary Tables 3-5)**. The Upset plot in **Figure 3A (left)**, summarizes the size of each DEG as well as the intersection amongst the three patients. Although there was a significant overlap amongst the DEGs between the three TTC7A patients **(Figure 3A and Supplementary Table 6**), in general the effect sizes were small. P1 and P3 appeared to share the most DEGs, however, many were in the opposing directions (n= 70 genes, BH= 12.95).

**Figure 3.**
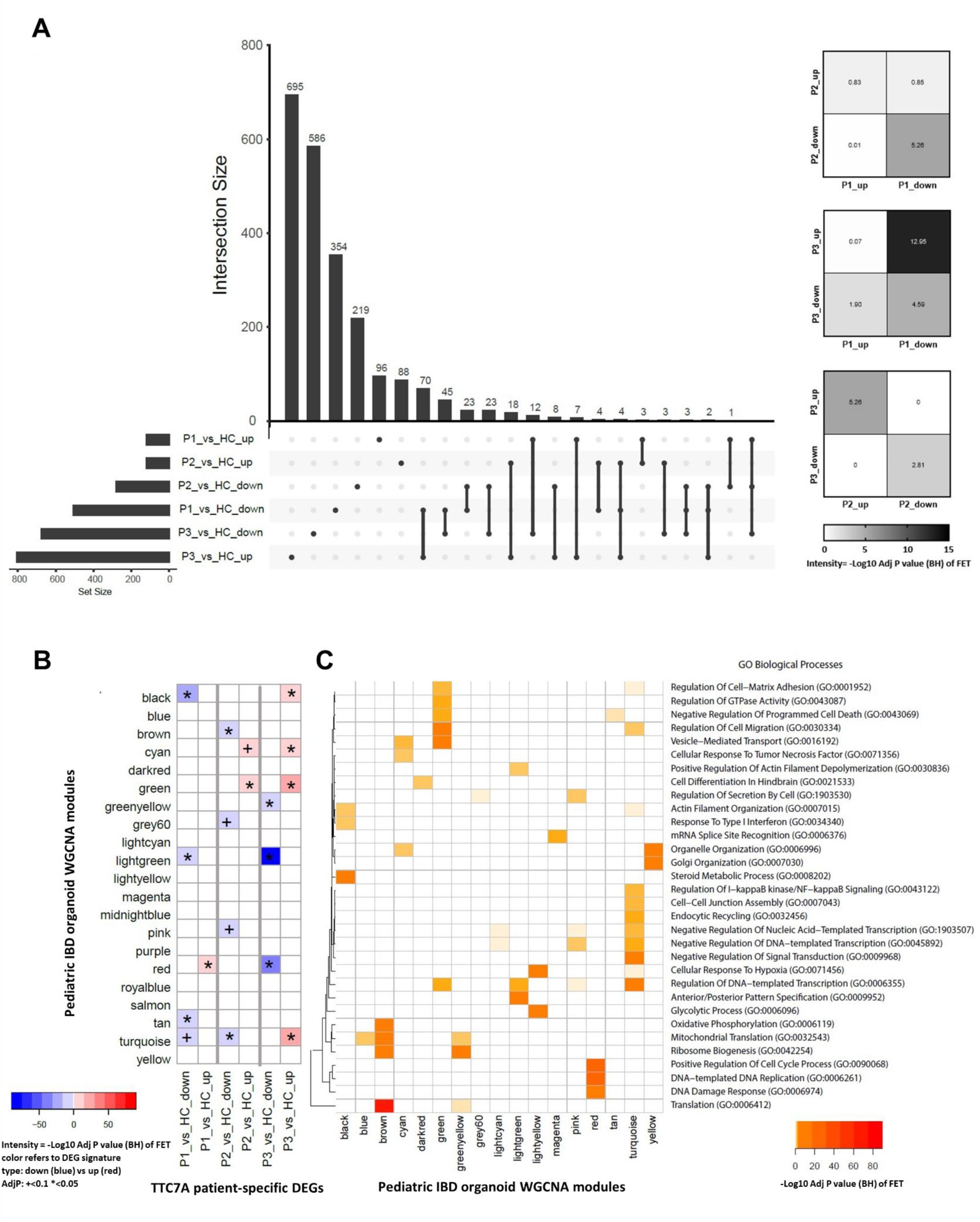
Intestinal organoids from patients with distinct *TTC7A* mutations reveal transcriptional heterogeneity. **(A)** Genes in organoids were defined as differentially expressed (DEGs) between each patient (P1, P2 or P3) versus healthy control (HC) (at P<0.05). The Upset plot (left) shows the overlap of the genes and the set sizes. The heatmap (right) shows the neg log10 Adj P value (BH) of the test of enrichment amongst the different DEGs. **(B)** The heatmap summarizes the neg log10 Adj P value (BH) for the enrichment of the DEGs (as described in A) in the various coexpression network modules (referred to by color name) as defined by WGCNA of a cohort of pediatric IBD and HC intestinal organoid transcriptomes (see methods). **(C)** Pathways associated with modules enriched in molecular alterations in TTC7A patient organoids. A heatmap summarizing the neg log Adj P value (BH) for the enrichment of GO biological processes in the WGCNA modules. Only a selected set of GO terms and modules are shown with full data available for GO BP and Reactome pathways in Supplementary Table 9 and 10.

Given the restricted number of technical replicates (N=2) for each TTC7A patient, we decided to pursue an alternative, more powerful strategy to identify potentially relevant biological pathways affected by the *TTC7A* variants. We projected each patient DEG (as up or down-regulated gene set) onto a WGCNA created from a larger cohort of HC and pediatric IBD intestinal organoid samples, including the *TTC7A* variant samples **(Supplementary Tables 1 and 2)**. The signed organoid WGCNA consisted of 22 modules of varying sizes as summarized in **Supplementary Table 7**. The modules significantly enriched at AdjP<0.1 **(Supplementary Table 8)** are shown in the heatmap in **Figure 3B**. Consistent with the poor overlap of the different TTC7A patient DEGs, in general, modules were found differentially enriched for by the various DEGs, supporting the idea that different mechanisms may underlie the phenotypic differences. For example, the green module was more significantly enriched for up-regulated genes in P2 (FE=3.2, AdjP=1.3×10^-6^) and P3 (FE=2.3, AdjP=5.3×10^-18^), than in P1 (FE=1.6, AdjP=0.33).

Consistent with P1 and P3 sharing the most opposing DEGs, the black module was significantly enriched for down-regulated genes in P1 (FE=3.4, AdjP=6.7×10^-18^) but up-regulated genes in P3 (FE=2.6, AdjP=1.8×10^-14^). Moreover, the red module was found significantly enriched for genes up-regulated in P1 (FE=2.3, AdjP=0.01) but down-regulated in P3 (FE=3.8, AdjP= 2×10^-43^).

As modules of coexpressed genes imply a shared co-function, we next evaluated the modules for pathway enrichments according to the GO biological processes (BP) and Reactome databases. A selection of GO BP terms found significantly enriched in the various modules is shown in the heatmap in **Figure 3C** with the full data available for GO and Reactome in **Supplementary Tables 9and 10**.

The pathways associated with the green module highlighted ‘regulation of GTPase Activity’; ‘negative regulation of programmed cell death’; ‘vesicle-mediated transport’; ‘trans-golgi network vesical budding’ and ‘RIPK1-mediated regulated necrosis’ to highlight a few. In contrast the black module highlighted terms associated with ‘metabolism’; ‘actin-filament organization’; ‘tight-junction’ and ‘interferon signaling’.

Thus, the black and green module appear to highlight different pathways associated with membrane organization and dynamics, as well as apoptotic pathways. Reduced expression of the black module in P1 and increased expression of the green module in P2 may help explain the heterogeneity observed in P1 and P2 phenotypes **(Supplementary Figure 5A)**.

The red module stands out for its notable enrichment in cell cycle processes and comprises genes that are down-regulated in P3 organoids **(Supplementary Figure 5A)**. This correlation potentially elucidates the observed hindered growth in P3 organoids. Finally, a striking enrichment of genes down-regulated in P3 and somewhat in P1 (but not P2) organoids was observed in the lightgreen module. This module was significantly enriched in genes associated with anterior/posterior pattern specification as well as embryonic organ morphogenesis and included many genes of the *HOXB*, *HOXA* family as well as *CDX1* **(Supplementary table 9)**. Notably, this effect was observed independent of the intestinal location (small vs large intestine) of the organoid **(Supplementary Figure 5B)**. This biology suggests that the fate determination of P3 organoids (and somewhat P1) may be dysregulated and potentially explain the phenotypic abnormalities observed. Interestingly, TTC7A has been reported to have functions in the nucleus, in chromatin organization and gene regulation. Consensus motifs found to potentially be bound by TTC7A included HOXA2 as well as EGR1, important transcriptional regulators of cell identity.^39^ This finding is particularly intriguing because it suggests a potential mechanism for the observed dysregulation of fate determination: if TTC7A directly regulates HOX genes and other developmental transcription factors, then mutations in *TTC7A* could disrupt this regulation, as observed in P1 and P3 organoids.

In summary, our investigation into the transcriptional profiles of intestinal epithelial organoids from three patients with distinct *TTC7A* variants revealed a limited concurrence among their DEGs. Further exploration via WGCNA identified distinct modules characterized by divergent biological pathways, each differentially enriched by the DEGs from the patients. This intricate pattern suggests the involvement of varied mechanisms underlying the phenotypic variations observed among these individuals and their intestinal organoids.

## Discussion

TTC7A deficiency is associated with a broad spectrum of intestinal and immune diseases including but not limited to VEOIBD and MIA with or without immunodeficiency of variable severity.^1,3,4^ Here we present the cases of three patients with distinct, deleterious heterozygous missense mutations in *TTC7A*, all diagnosed with early-onset IBD, two of whom also exhibited features of CID. However, despite sharing these diagnoses, the patients exhibited distinct clinical features, varying disease severity, and differing treatment requirements, for both their intestinal and immunological conditions. Notably, CIPO was previously reported in a patient as a novel phenotype associated with TTC7A,^2^ marking P1 the second documented case of a TTC7A patient displaying CIPO. We hypothesize that part of the diversity in clinical presentation stems from the presence of distinct disease-causing *TTC7A* variants among these patients.

In support of our hypothesis, we also observed striking diversity in the epithelial characteristics of the patient-derived intestinal organoids. Organoid lines varied in growth patterns, morphologies, apicobasal polarities, levels of TTC7A protein expression, and drug responses. Notably, ROCK inhibitor enhanced morphology, growth, and viability in both P2 and control duodenal organoids, but not P1. This suggests ROCK inhibition may improve duodenal epithelial health independently of TTC7A dysfunction, contradicting previous reports.^11,19^ Leflunomide, previously shown to improve TTC7A deficiency by reducing caspase-dependent apoptosis in vitro, failed to restore normal polarization in P2 organoids, likely due to the absence of elevated caspase activity.^23^ Similarly, it had no clinical benefit in a patient with neonatal-onset IBD carrying two *TTC7A* variants: a pathogenic nonsense mutation in exon 9 (E215X) and a variant of uncertain significance in exon 19 (c.2355+4A>G).^40^ Furthermore, RNA-seq analysis revealed different transcriptional profiles between these patient organoids, highlighting the diverse biological mechanisms underlying TTC7A variant-driven heterogeneity.

Moreover, compared to previous studies of *TTC7A*-mutant organoids, our patient-derived organoids with different *TTC7A* mutations exhibited distinct epithelial phenotypes. Notably, in contrast to prior findings on colon organoids from a patient with biallelic missense mutations,^23^ P1 ileal organoids maintained an intact epithelial transport function. Additionally, P1 organoids retained normal apicobasal polarity which contrasts with the disrupted polarity previously described in ileal and rectal organoids from patients with distinct homozygous mutations.^11,19^ Unlike earlier observations of elevated caspase-3 immunostaining, we did not detect increased caspases 3 and 7 activity in P1 or P2 organoids.^19^ Our intestinal organoid model highlights the significant impact of diverse mutations within *TTC7A* on epithelial phenotypes and drug responses, illustrating the complex relationship between genetic variants and therapeutic outcomes.

In a previous study using TTC7A^-/-^ Caco cells, we individually characterized the molecular and cellular effects of different *TTC7A* missense mutations, including K606R (in P1), G173V (in P2), E343K (in P3), and L802P (in P3), and classified them based on their severity of impact. K606R and G173V were deemed pathogenic due to their association with aberrant transcriptional signatures, whereas E343K and L802P were classified as benign or inconclusive, as they did not significantly alter the epithelial phenotype.^7^

The K606R and S672P variants, identified in cis on the same TTC7A allele in P1, have likewise been reported in cis in six other patients. Their identical low allele frequency in gnomAD (0.0022 across all genomes) supports a shared origin on a rare ancestral haplotype. In a cohort of 401 pediatric patients who underwent whole-exome sequencing (WES), five individuals were found to carry both variants on the same parental chromosome (either maternally or paternally inherited). Four of these had a synonymous variant on the second *TTC7A* allele, and one had no additional *TTC7A* mutation, mirroring the configuration observed in P1. All five were diagnosed with pediatric IBD, with variable clinical presentations: four had CD, only one of whom required surgery for stricturing disease, and one had UC. Notably, the transmitting parents were unaffected. In this cohort, in silico gene-level burden analysis, performed on an individual basis by aggregating deleteriousness scores for all exonic variants within TTC7A, did not implicate *TTC7A* variation as a significant contributor to disease in the IBD cohort.^41^ These findings suggest that the presence of K606R and S672P in cis, such as observed in P1, may be insufficient on their own to cause disease, and that additional genetic or environmental modifiers may influence penetrance and clinical expression. In another study of a patient with MIA/CID, both K606R and S672P were found on the maternal allele, along with a 4-nucleotide deletion on the paternal allele causing Exon 7 skipping.^4^ In contrast, two other patients, one with common variable immunodeficiency/enteropathy, and another with Severe Combined Immunodeficiency /MIA, carried K606R and S672P in trans, each inherited from a different parent, representing a compound heterozygous state.^42,43^

Although TTC7A deficiency is typically considered an autosomal recessive condition, other monoallelic *TTC7A* mutations have been reported. These include a deletion of exons 2 and 3 associated with intestinal atresias/CID,^4^ and a missense variant (L823P) in exon 20 linked to IBD/CID in one patient and delayed-onset CID in another.^44,45^ Furthermore, single-allele mutations have been observed in other monogenic IBD genes typically associated with autosomal recessive inheritance, such as *HPS1*, *FERMT1*, *LRBA*, *MASP*, *NCF4*, *SLC37A4*, and *SLC9A3.*^46^ These findings illustrate our incomplete understanding of *TTC7A* genetics. For example, some mutations might have a dominant negative effect that cannot be compensated by the function of the normal protein produced from the healthy allele. This could explain why some individuals, like P1, with only one mutated allele still develop disease. Beyond these reported cases of monoallelic mutations, another possibility is that P1 harbors undetected variants. Given that WES often misses copy number variants, intronic variants, synonymous variants, and inversions, and provides incomplete coverage of certain regions in known monogenic IBD genes,^47–50^ it is possible that P1, with apparent monoallelic inheritance, harbors undetected variants, consistent with an autosomal recessive pattern.

Despite having the mildest intestinal disease, P2 organoids, with heterozygous G173V and L452P mutations, exhibited a significantly abnormal phenotype, disrupted polarity and reduced TTC7A protein, while maintaining normal morphology, growth, and viability. It remains unclear which test best correlates with IBD severity. While the abnormal phenotype in P2 organoids supports the classification of G173V as pathogenic, this variant alone did not previously show TTC7A protein downregulation, as observed in P2 organoids.^7^ This suggests that TTC7A downregulation may be attributed to the L452P variant. While consistently predicted to be pathogenic by computational models, L452 has no prior record in existing documentation. Further functional studies are needed to clarify its specific effects.

Although both mutations identified in P3 (E343K and L802P) were classified as benign or inconclusive in our previous study, P3 presents with severe intestinal disease.^7^ Computational prediction tools consistently classified both L802P and E343K as pathogenic, with the exception of MutationTaster 2021, which predicted E343K to be benign. Notably, the E343K variant was previously identified in a patient homozygous for E343K who had CID but no intestinal disorder.^5^

The discrepancies between our earlier findings on the individual effects of these variants and the clinical phenotypes observed in patients, and in patient-derived intestinal organoids may be explained by several factors. These include intrinsic differences between Caco-2 cells and more physiologically relevant organoid models, as well as patient-specific genetic background, environmental influences, and epigenetic modifications. Additionally, while each variant may appear functionally mild on its own, their combined presence in compound heterozygosity, as observed in our patients, may lead to synergistic or additive defects not detectable in simplified models. It is also possible that certain variants exert context-dependent effects, manifesting differently across tissues or disease states.

These findings underscore the need for more physiologically relevant platforms, such as intestinal organoids that encompass all differentiated epithelial lineages. Using CRISPR-Cas9–based technologies including base editing and prime editing, which are particularly well-suited for introducing or correcting missense mutations without generating double-strand breaks, these models can be precisely engineered to carry *TTC7A* variants.^51,52^ This allows for systematic assessment of variant effects both individually (in a homozygous context) and in combination (as compound heterozygotes). When coupled with advanced analytical tools, such as single-cell RNA sequencing and integrative multi-omics profiling, these systems offer a powerful approach for dissecting variant-specific pathogenic mechanisms and informing personalized therapeutic strategies.

## Conclusion

Distinctive pathogenic mutations in *TTC7A* have been shown to give rise to varying clinical presentations, heterogeneous cellular phenotypes, differences in drug responses and diverse transcriptional signatures in intestinal organoid models. This highlights the varying severity of the consequences of individual *TTC7A* mutations and proposes that multiple mechanisms may contribute to TTC7A-related disorders. Further research is needed to fully dissect the molecular consequences of individual *TTC7A* mutations. The heterogeneity in molecular ramifications and resulting clinical presentations may have significant implications when outlining therapeutic interventions for individuals with certain *TTC7A* mutations.

## Acknowledgements

We thank all the patients and their families who consented and participated in this study as part of the Helmsley VEOIBD (www.VEOIBD.org) consortia and the STEM study. We thank Utrecht Sequencing Facility (USEQ) for performing RNA sequencing. We thank HUB organoids for generating intestinal organoids derived from P2 biopsies.

## Author contributions

EN and MM contributed to study and experiment designs. ZS, PD, MV, and JB performed the experiments. CA designed and performed RNA-seq analysis and interpretation. ZS performed data analysis and wrote the manuscript. CL, SS, CK, SK, LC, and DK provided patient biopsies and clinical data. JMB provided resources for FIS assay. EN, EK, CL, SK, and LC provided intellectual content. EN and EK edited the manuscript. All of the authors fulfill the criteria for authorship and read and approved the final manuscript.

## Data availability statement

The processed RNA sequencing data will be made available upon publication.

