## Supplement for "Distinct heterozygous *TTC7A* missense variants lead to different intestinal epithelial phenotypes in pediatric IBD"

### Supplementary Materials

**Supplementary Table 1.** Pediatric IBD organoid RNA-seq cohort

| No. | Sample | Donor | Batch | Location | Gender | Age_at_sample collection |
|---|---|---|---|---|---|---|
| 1 | MV18Cae0H | D018 | E | caecum | F | 19 |
| 2 | MV48Cae0H | D048 | E | caecum | F | 11 |
| 3 | MV50Cae0H | D050 | E | caecum | F | 6 |
| 4 | VEO90H | D009 | A | caecum | M | 5 |
| 5 | X12Cae0H | D012 | D | caecum | F | 2 |
| 6 | X7Cae0H | D007 | D | caecum | F | 5 |
| 7 | X9Cae0H | D009 | D | caecum | M | 5 |
| 8 | MdV17C0H | D017 | C | colon | F | 2 |
| 9 | MdV37C0H | D037 | B | colon | F | 13 |
| 10 | MdV3C0H | D003 | B | colon | M | 2 |
| 11 | MV52Sig0H | D052 | E | colon | M | 2 |
| **12** | **MV65.0H** | **D065 (P3)** | **E** | **colon** | **M** | **4** |
| **13** | **VEO65C0H** | **D065 (P3)** | **H** | **colon** | **M** | **4** |
| 14 | MV74.0H | D074 | E | colon | M | 7 |
| 15 | X14Sig0H | D014 | D | colon | M | 3 |
| 16 | X17Asc0H | D017 | D | colon | F | 2 |
| 17 | X85Sig0H | D085 | G | colon | M | 5 |
| 18 | X90Des0H | D090 | G | colon | M | 4 |
| 19 | X95TC0H | D095 | F | colon | M | 10 |
| 20 | X97TC0H | D097 | F | colon | F | 1 |
| 21 | X98AC0H | D098 | F | colon | M | 3 |
| 22 | MdV11D0H | D011 | C | duodenum | F | 2 |
| **23** | **STE165Du0H** | **STE165 (P2)** | **I** | **duodenum** | **M** | **13** |
| **24** | **STE165duo0H** | **STE165 (P2)** | **F** | **duodenum** | **M** | **13** |
| 25 | X12Duo0H | D012 | G | duodenum | F | 2 |
| 26 | MdV11I0H | D011 | B | ileum | F | 2 |
| 27 | MdV14I0H | D014 | B | ileum | M | 3 |
| 28 | MdV29I0H | D029 | B | ileum | F | 8 |
| 29 | MdV39I0H | D039 | B | ileum | M | 6 |
| 30 | MV39Cae0H | D039 | E | ileum | M | 6 |
| **31** | **MV67ile0H** | **D067 (P1)** | **E** | **ileum** | **M** | **18** |
| **32** | **VEO67il0H** | **D067 (P1)** | **H** | **ileum** | **M** | **18** |
| 33 | VEO140H | D014 | A | ileum | M | 3 |
| 34 | X1ile0H | D001 | G | ileum | M | 5 |
| 35 | X12Rec0H | D012 | G | rectum | F | 2 |
| 36 | X3Rect0H | D003 | D | rectum | M | 4 |
| 37 | X45Rect0H | D045 | D | rectum | M | 6 |
| 38 | X51Rec0H | D051 | F | rectum | F | 8 |
| 39 | X7Rect0H | D007 | D | rectum | F | 5 |

**Supplementary Table 2.** Control organoid RNA-seq cohort

| No. | Sample | Donor | Batch | Location | Gender | Age_at_sample collection |
|---|---|---|---|---|---|---|
| 1 | MV36Cae0H | D036 | E | caecum | M | 1 |
| 2 | X26Cae0H | D026 | F | caecum | M | 7 |
| 3 | MdV13C0H | D013 | C | colon | M | 3 |
| 4 | MdV32C0H | D032 | C | colon | M | 3 |
| 5 | MdV33C0H | D033 | B | colon | M | 2 |
| 6 | MdV34C0H | D034 | C | colon | M | 3 |
| 7 | MdV35C0H | D035 | B | colon | F | 3 |
| 8 | MdV44C0H | D044 | C | colon | M | 5 |
| 9 | VEO130H | D013 | A | colon | M | 3 |
| 10 | VEO32C0H | D032 | H | colon | M | 3 |
| 11 | VEO34C0H | D034 | H | colon | M | 3 |
| 12 | X8Asc0H | D008 | D | colon | M | 13 |
| 13 | X8Sig0H | D008 | D | colon | M | 13 |
| 14 | X91Des0H | D091 | G | colon | M | 13 |
| 15 | X94Sig0H | D094 | F | colon | F | 11 |
| 16 | STE76Du0H | STE076 | I | duodenum | F | 8 |
| 17 | HC4il0H | HC4 | I | ileum | M | 1 |
| 18 | MdV26I0H | D026 | B | ileum | M | 7 |
| 19 | MdV28I0H | D028 | B | ileum | F | 3 |
| 20 | MdV43AI0H | D043 | B | ileum | M | 6 |
| 21 | STE150il0H | STE150 | H | ileum | F | 41 |
| 22 | STE94il0H | STE094 | I | ileum | M | 12 |

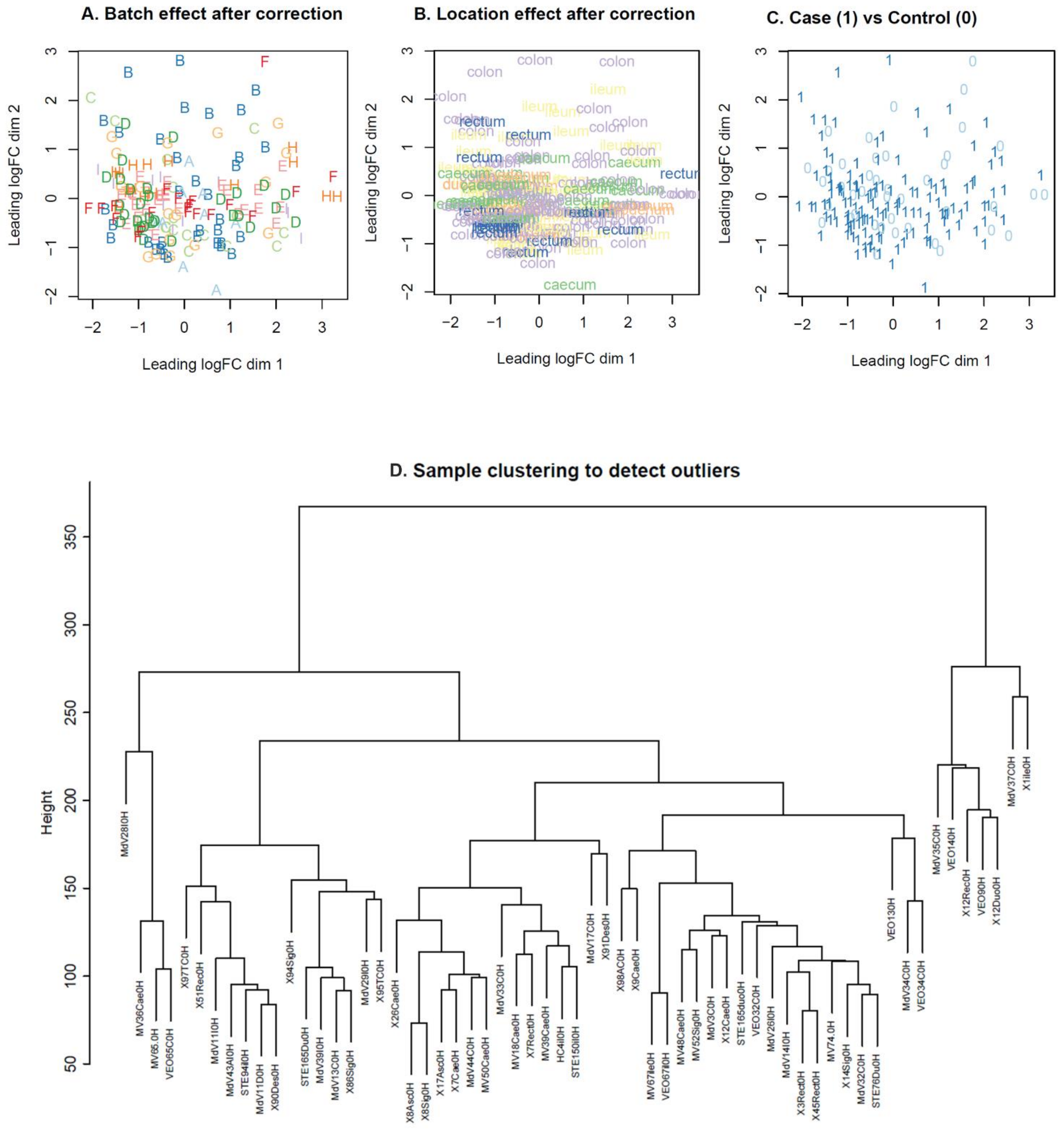


**Supplementary Figure 1. Quality control plots related to the cohort of intestinal organoid transcriptomes**. **A-C**, Multidimensional scaling (MDS) plots to visualize the effect of Batch **(A)**, intestinal region **(B)** and Case/control status **(C)** on expression. The dimensions represent the underlying factors that explain the dissimilarities among the observations with the clusters representing the groups of observations that are similar to each other. **(D)** A dendrogram showing the results of hierarchical clustering (method ='ward') of the transcriptomes of 61 intestinal organoid samples in order to detect potential outlier samples.

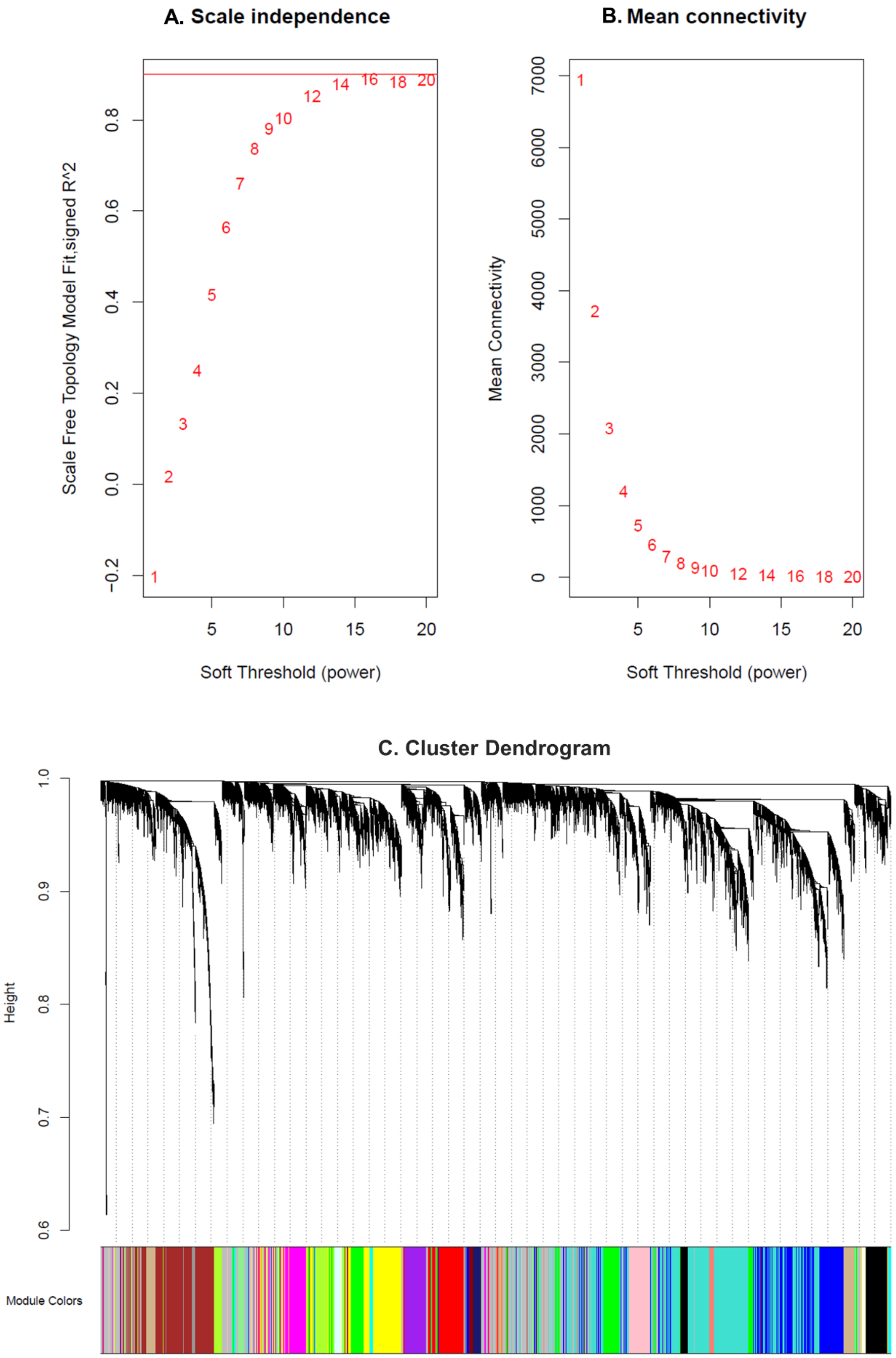


**Supplementary Figure 2. (A)** Scale independence, **(B)** mean connectivity plots and **(C)** cluster dendrogram for the signed WGCNA analysis of cohort of pediatric IBD and HC intestinal organoid transcriptomes ( see methods). A power threshold of 12 was chosen for the network generation. Each dot represents a sample.

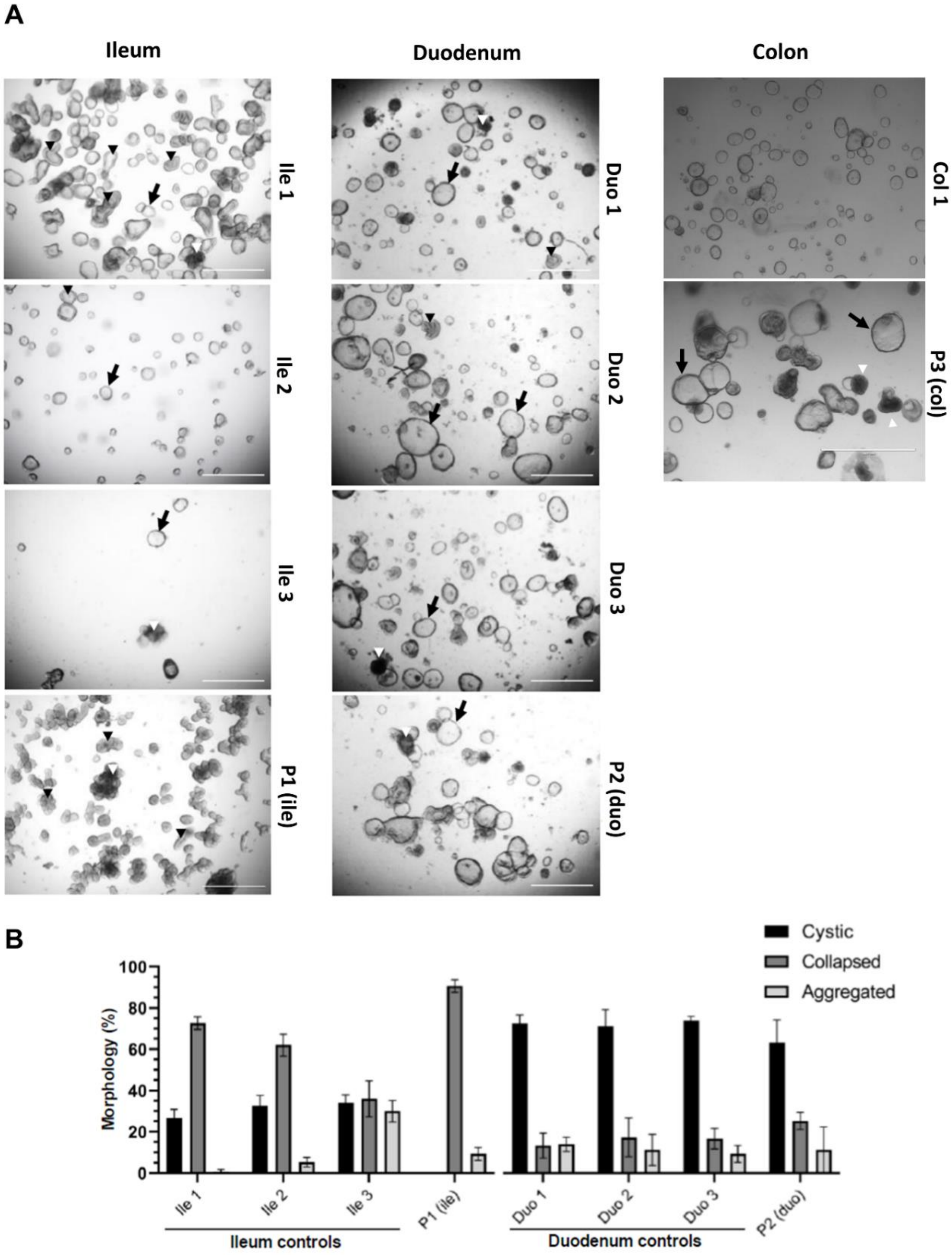


**Supplementary Figure 3. Morphological phenotypes of intestinal organoids**

**(A)** Organoids were seeded by mechanical shearing and overall organoid condition was assessed for three parameters: circular or swollen organoids were considered normal and classified as 'cystic' (black arrow), cell-dense, non-cystic structures were deemed 'collapsed' (black arrowhead) and darkened aggregations of organoid tissue where distinct shapes could not be identified were assigned an 'aggregated' phenotype (white arrowhead). P1 (ile), P2 (duo) and P3 (col) represent different TTC7A patient-derived organoids from ileum, duodenum, and colon respectively. Ile 1-3 (ileum), Duo 1-3 (duodenum) and col 1 (Colon) are healthy control lines for P1 (ile), P2 (duo), and P3 (col) respectively. Representative images were obtained with a stereomicroscope (EVOS) at a magnification of 4x. **(B)** Organoids were seeded by mechanical dissociation and morphology was quantified by assessing 50 organoids per field (N=3) . Images were obtained with a stereomicroscope (EVOS) at a magnification of 4x. (Mean ± SD).

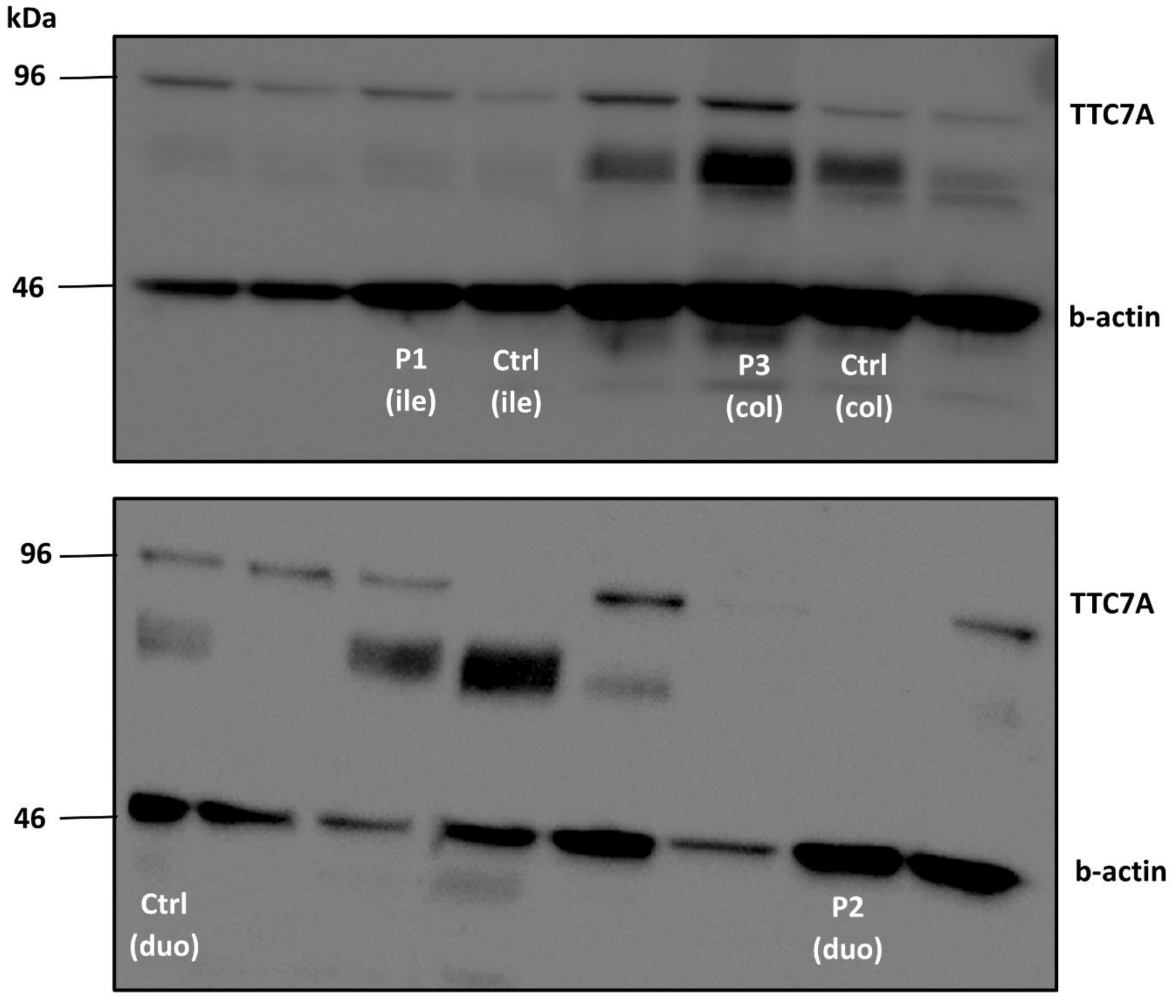


**Supplementary Figure 4. Original Western Blot Images**

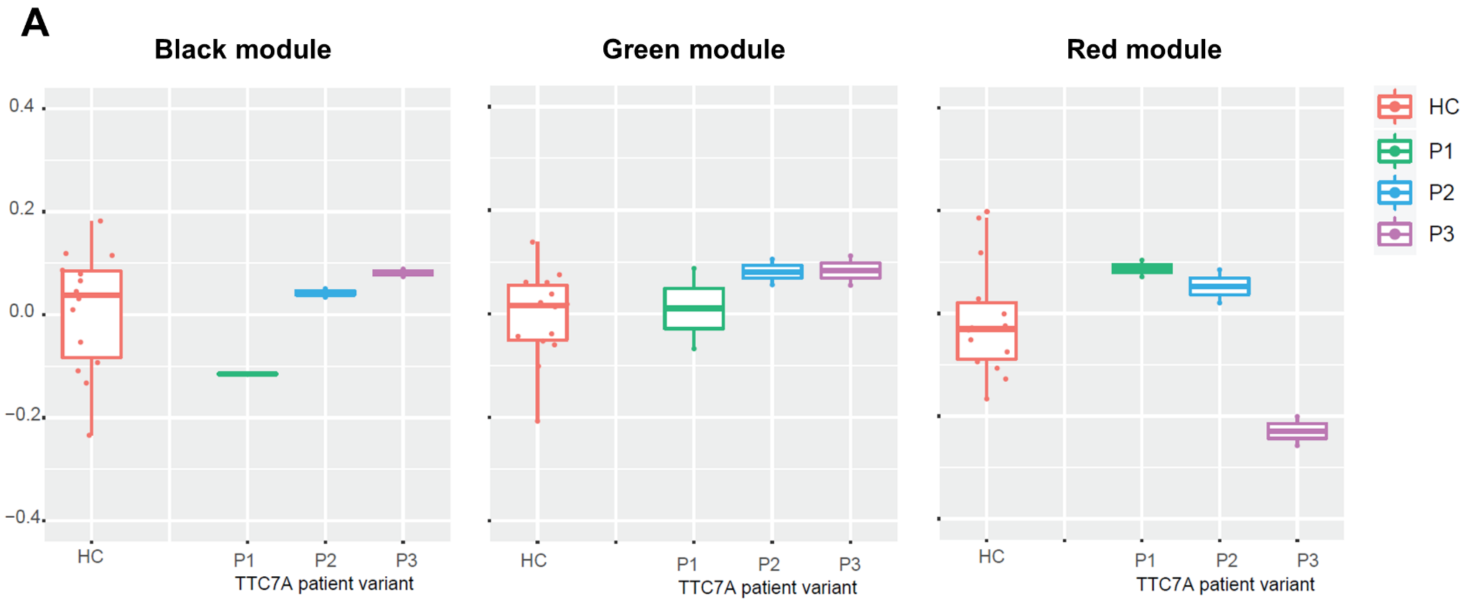


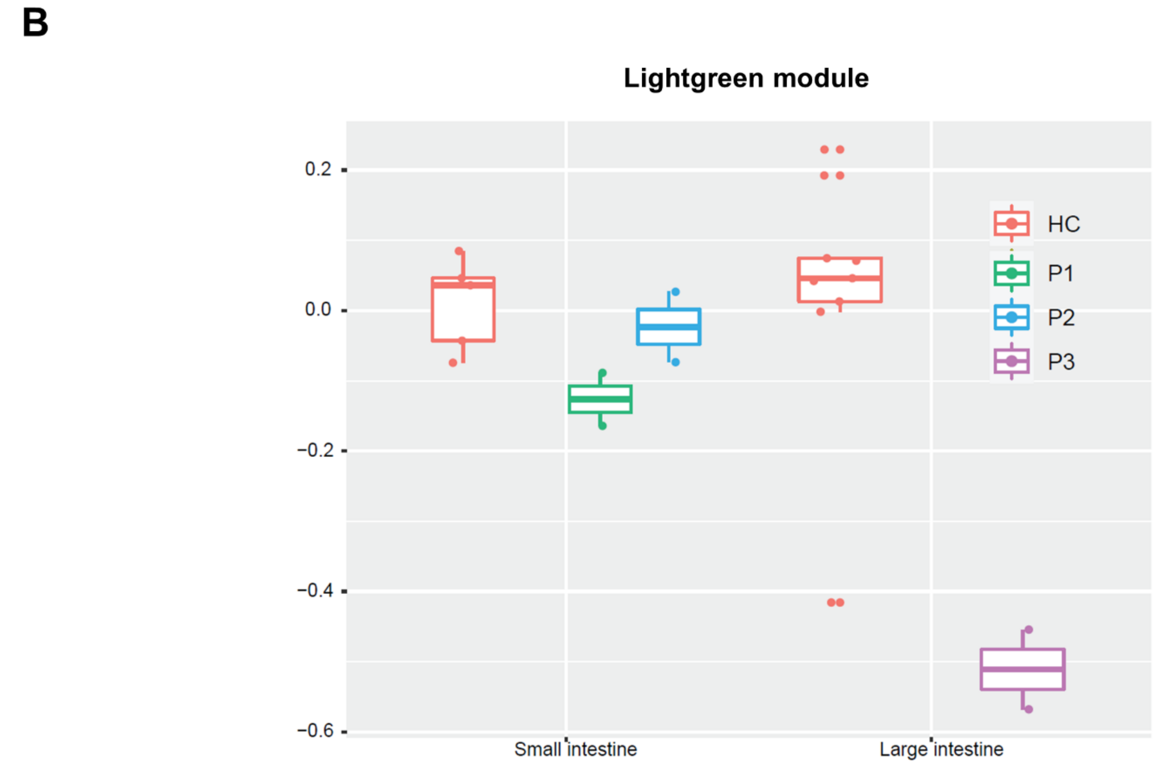


**Supplementary Figure 5. (A-B)** Boxplots summarizing expression of various WGCNA module eigengenes within healthy controls (HC) or the various TTC7A patient variants. **(B)** Module eigengene expression is further segregated by sampling location (small vs large intestine) for the various patient organoids. Box plots were utilized to demonstrate variance in HC's, even though patient samples only had duplicate values.
